# Multiple patterns of selectivity in superior colliculus control visual search

**DOI:** 10.1101/2025.11.11.687826

**Authors:** Abe Leite, Hossein Adeli, Robert M. McPeek, Gregory J. Zelinsky

## Abstract

Visual search is a ubiquitous behavior for many animal species, but efficient search performance requires a relatively complex integration of target guidance and behavioral history signals. In this study, we investigate how the superior colliculus (SC), a midbrain area implicated in the control of overt spatial attention, integrates these two control signals in a multi-target search task. Monkeys made sequences of saccades to stimuli presented in a grid while searching for a “true” (reward eliciting) target amid color-coded targets and distractors. The efficient performance of this task requires integrating a target guidance signal with a history of previously fixated locations in order to guide search to new potential targets that might offer reward. Theories of systems neuroscience suggest that this multi-factor search behavior might be controlled using either mixed selectivity, dynamic selectivity, or pure selectivity. We find evidence for SC neurons exhibiting each of these patterns and conclude that this brain structure participates in circuits representing target similarity and previously fixated locations, in addition to a circuit responsible for integrating these signals using the mechanisms of dynamic and mixed selectivity. We also introduce a novel time-series partial information decomposition analysis that provides a rich and direct view of how a neuron allocates its representational capacity. These findings support an emerging view of the superior colliculus as much more than a bottom-up priority map, but instead an important site for the flexible cognitive control of visually-guided behavior.

## 1. Introduction

Visual search is ubiquitous in animal behavior – organisms from fish to humans commonly look for visual objects in their environments to mediate the everyday demands of existence. Search is also a complex behavior requiring the flexible integration of multiple factors. Two of the most important factors for search efficiency are top-down target guidance and a history of previously-searched locations (Wolfe and Horowitz, 2017). Target guidance preferentially directs attention to visual features matching the target, and a history of previously searched locations prevents attention from returning to locations that were already determined not to have a target.

This study focuses on how the factors of target guidance and search history are represented by neurons in the primate superior colliculus (SC) as a monkey is actively engaged in a visual search task. The SC (optic tectum in non-mammals) is a highly-conserved midbrain area that acts as the primary center of visuospatial cognition and navigation behavior in non-mammalian vertebrates (Cooper and McPeek 2021). In mammals, the SC integrates sensory and cognitive information from multiple brain areas (Liu et al., 2022), and it plays a critical role in visually-guided behavior, particularly eye movements and covert attention (Sparks and Hartwich-Young, 1989; Gandhi and Katnani 2011; Krauzlis, Lovejoy, Zénon 2013; Basso and May 2017). In recent years, a consensus has emerged that a primary function of SC is to help instantiate a priority map for saccade target selection (e.g., McPeek and Keller 2002; Krauzlis et al. 2004; Fecteau and Munoz 2006; Kim and Basso 2008; Shen et al. 2011; Song et al. 2011; Mysore and Knudsen 2011). Thus, the primate SC is an ideal brain structure in which to study how task variables are encoded and combined in a priority computation for target selection.

### 1.1 Multi-target grid search task

Our study uses behavior and neural data that were collected from monkeys performing a task in which they made sequences of saccadic eye movements to search a grid of stimuli (Conroy, Nanjappa, and McPeek, 2023). Monkeys are presented with an array of 15-25 colored disk stimuli arranged in a grid (Figure 1A). Roughly one third of disks have the target color, either red or green depending on trial block, and the remaining two thirds are distractor disks having the non-target color. While all the target-colored discs have an identical visual appearance, only one of the disks is the *true target*, meaning the disk whose fixation results in termination of the trial and receipt of a reward. Efficient search behavior would therefore be the serial fixation of each target-colored disk while avoiding previously searched disks and distractors, in order to find the true target as quickly as possible.

**Figure 1.**
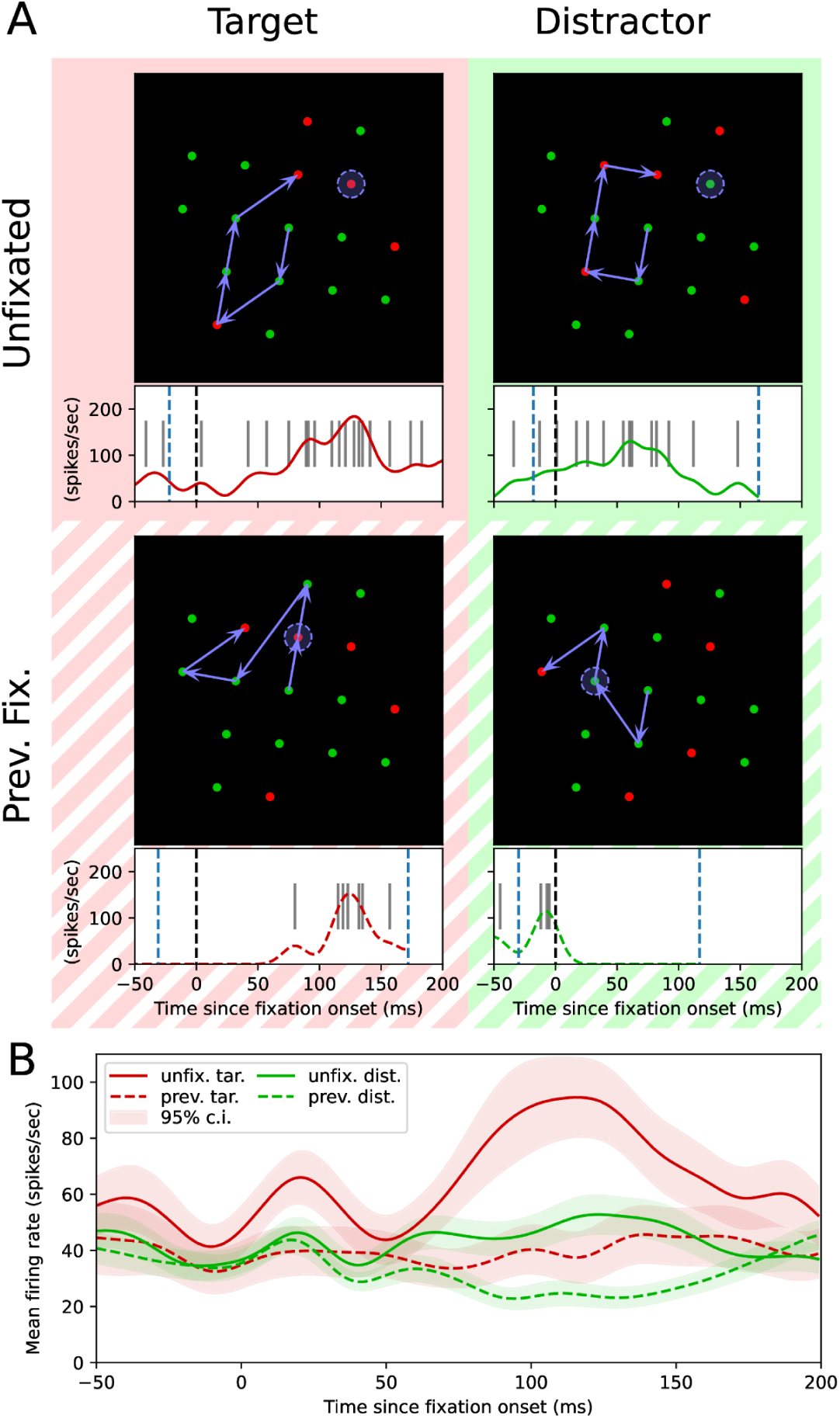
This recorded neuron’s response depends on the properties of the stimulus in its receptive field (RF). (A) Sample displays and neural responses under the four combinations of our two factors: whether the disk in the neuron’s RF (indicated with a dashed blue circle) was of the target or distractor color and whether it was previously fixated or not. In each display, the target color is red and the monkey’s eye movements are shown by blue arrows (starting from the central fixation point). Below each display is the response of the recorded neuron. Firing rate is shown over the course of the fixation which brought the neuron’s RF to its indicated location. Fixation onset is indicated by the dashed black line; preceding and following saccade onsets are shown with dashed blue lines. A raster plot of spikes is indicated by the thin black lines. (B) The neuron’s response grouped by the four stimulus conditions and averaged over multiple trials. Shaded ranges indicate a 95% confidence interval. This neuron’s response was greatest when the disk in its receptive field shared the target color and had not been previously fixated.

To control this efficient selection of search targets, the SC must integrate two pieces of information about each disk: whether it shares the **target color** and whether it has been **previously fixated** during the current search trial. We contrast previously-fixated disks with disks that have yet to be fixated in the search, and we use the term “unfixated” to refer to disks that had never been fixated before the current point in the search trial (rather than to disks that are not currently fixated). These target-color and previously-fixated properties map onto well-studied cognitive processes: feature-based attention guidance to search targets (Treisman and Gelade, 1980; Zelinsky and Sheinberg, 1995) and the inhibitory tagging of previously attended locations (Klein and Macinnes, 1999); this lends broad theoretical importance to the question of how these factors are encoded in the brain.

Figure 1A shows behavioral and neural data on disk stimuli in each target color and previous fixation condition. For each trial, a grid array was generated such that when fixating one grid element, a neighboring element would fall into the receptive field (RF) of the neuron under study (with the exception of fixations of some elements at the edge of the grid). The search-relevant properties of the stimulus in the neuron’s RF varied, resulting in four conditions corresponding to specific combinations of properties. In the *target color / previously unfixated* condition (Fig. 1A; upper left), the disk in the neuron’s RF has the target color and has not yet been fixated. This is the condition when search is most expected to be guided to the recorded disk, and the enhanced neural response for the illustrated neuron is consistent with a target guidance signal. The *target color / previously fixated* condition (Fig. 1A; lower left) best captures when search is expected to be influenced by inhibitory tagging; the disk in the neuron’s RF has the target color but has been previously fixated. The right two panels in Figure 1A illustrate conditions in which a disk having a non-target color appears in the neuron’s RF, with and without previous fixation. Neural firing rate data were analyzed over a −50ms to +200ms window around each fixation onset. Figure 1B plots the mean firing rate for this neuron during a fixation, grouped by the four conditions, replicating Conroy et al.’s analysis (2023). That analysis found that the population responded most strongly to unfixated targets, and that 52 of the 113 neurons carried a significant previous fixation signal. Our goal in this paper is to reappraise these data to address the broader question of which computations SC uses to support flexible search behavior.

### 1.2 Theories of neural selectivity

Understanding the neural computations underpinning the flexible integration of multiple sources of information is a grand challenge in contemporary neuroscience, and a broad range of theoretical approaches have risen to meet it. Here, we focus on three families of models for the role a single neuron might play in this integration: (1) mixed selectivity, (2) dynamic selectivity, and (3) pure selectivity. These theories of selectivity rely on a set of experimenter-defined *fundamental task variables* that are assumed to represent the basic factors supporting a decision in a task. In the multi-target search task, the fundamental variables supporting saccade target selection are whether a disk shares the target color and whether it has been previously fixated. Our study focuses on the role of SC neurons in integrating these two sources of information, which is a prerequisite to efficient search behavior.

#### 1.2.1 Pure selectivity

One way for a population of neurons to represent multiple sources of task-relevant information is to have sub-groups of neurons become purely selective to the different information sources. While this form of pure selectivity provides a good explanation, for instance, of simple cells in V1, it has been rarely observed in neurons outside of low-level sensory and motor areas (Tye et al., 2024). Modeling work also finds purely-selective neurons to be the exception rather than the rule. Evidence for pure selectivity is clearest in those *in silico* experiments where a single, fundamental computation is shared across many different tasks (Yang, Joglekar, Song, Newsome, and Wang, 2019).

We test whether the SC contains purely-selective neurons representing the fundamental factors of target color and previous fixation in the search context. On the basis of modeling work, we consider it plausible that SC may contain neurons purely selective to previous fixation, as this factor is important to many visual tasks other than search and would seem to rise to the level of needing a circuit dedicated to its computation.

#### 1.2.2 Mixed selectivity

Mixed selectivity theory (Rigotti et al., 2013) models a neuron’s activation as being dependent on multiple fundamental task variables. This is typically assessed by showing that no single variable is sufficient to explain the neuron’s firing rate – that is, that a model incorporating multiple variables explains the neuron’s firing rate better than any model incorporating only a single variable.^1^

Among the accounts of why mixed selectivity is observed (see also Hirokawa, Vaughan, Masset, Ott, and Kepecs, 2019), nonlinear category-free selectivity considers a population of neurons as an information reservoir whose role is to support downstream and task-specific decisions. According to this variant of mixed selectivity theory, the predicted neural response should not merely depend on multiple variables, but should be best explained by a nonlinear combination of those variables (Kaufman et al., 2022). Further, selectivity should be category-free (Raposo, Kaufman, and Churchland, 2014), meaning that each neuron has a distinct response pattern, rather than belonging to a ‘category’ of neurons that share a response pattern. (The absence of categories is generally tested by assessing the clustering of neurons’ responses within a space defined by the fundamental task variables. Clustering implies categories of neurons, while little clustering indicates a category-free population of neurons.)

The properties of being nonlinear and category free are most powerful when they act in concert. If every neuron is well-explained as a *linear* combination of task variables, then even in the absence of categories of neurons, the rank of the population response space cannot exceed the number of task variables. However, if neurons encode distinct *nonlinear* combinations of the task variables, then every neuron contributes something new to the population’s representation space, without any limit on the dimensionality of the population representation. This is theoretically significant because a reservoir of such neurons maximizes the number of possible linear decisions that a downstream *readout* neuron supporting a particular computation could represent. This allows the rapid learning of new computations by such readout neurons.

Mixed selectivity has been observed in evolutionarily-young cortical and subcortical forebrain areas, such as area V6a (Diomedi, Vaccari, Filipini, Fattori, and Galletti, 2020), prefrontal cortex (Dang, Li, Pu, Qi, and Constantinidis, 2022), and the hippocampus (Ledergerber et al., 2021), but has not to our knowledge been reported in phylogenetically older midbrain areas such as SC.

#### 1.2.3 Dynamic selectivity

Neurons can also integrate the information from multiple variables by representing different information at different times. For instance, coarse category information is encoded more rapidly by IT neurons than fine category information, and the same neurons encode both levels of information (Sugase, Yamane, Ueno, and Kawano, 1999). More recently, this phenomenon has been explored in the context of face detection and recognition, where it is described as switching of the neural code (Shi, Bi, Hesse, Lanfranchi, Chen, and Tsao, 2023) or temporal multiplexing (She, Benna, Shi, Fusi, and Tsao, 2024). Here, we adopt the broader term *dynamic selectivity* to refer to cases where a neuron’s selectivity changes based on the temporal dynamics of the system it belongs to.

Consistent with Sugase et al. (1999)’s pioneering work, Shi et al. (2023) found that many face-selective neurons in macaque IT initially appear to be face *detectors* – responding equally to all faces – but after a “rapid, concerted switch” over a 100ms-120ms period from stimulus onset, these neurons change their response pattern to act as face *discriminators* – responding selectively to particular face features such as interocular distance. The authors’ theory is that each neuron first represents that a face is, in fact, present, and then represents discriminative details about the specific face being seen. Such selectivity is theoretically advantageous because it avoids neurons ‘wasting’ their representational resources by encoding combinations of factors that are irrelevant to the current decision. It can be thought of as a way of maximizing the neuron’s utility over multiple behaviors and multiple portions of a single behavior, without category-free selectivity’s disadvantage of forcing the neuron to become useful only through auxiliary readout neurons.

### 1.3 Partial information decomposition

To distinguish between the reviewed theories of neural selectivity, we apply an emerging information theoretic technique known as *partial information decomposition* (PID) (Williams and Beer, 2010). Rather than relying solely on mean firing rates, this tool analyzes firing rate *distributions* to quantify the information that a neuron encodes about multiple task variables. PID then decomposes this quantity into information that is **redundant** between multiple variables (explainable given any one of them), **unique** to a single variable (explainable given it but not given the others), and **synergistic** between multiple variables (explainable given all of them, and no other combination).^2^

Figure 2A shows the terms produced by a 2-variable PID analysis of the information encoded by a neuron’s firing rate in the context of multi-target search. Information about the task variables of target color and previous fixation can be decomposed in four ways: (1) redundant information that can be predicted given either factor, (2) unique information about target color, (3) unique information about previous fixation, and (4) synergistic information about the interaction between the factors. As the Venn diagram depicts, the mutual information between firing rate and target color is the sum of both redundant and unique components, as is the mutual information between firing rate and previous fixation. The mutual information between firing rate and the joint distribution of target color and previous fixation (the total explainable information encoded in the firing rate) is the sum of all four terms.

**Figure 2.**
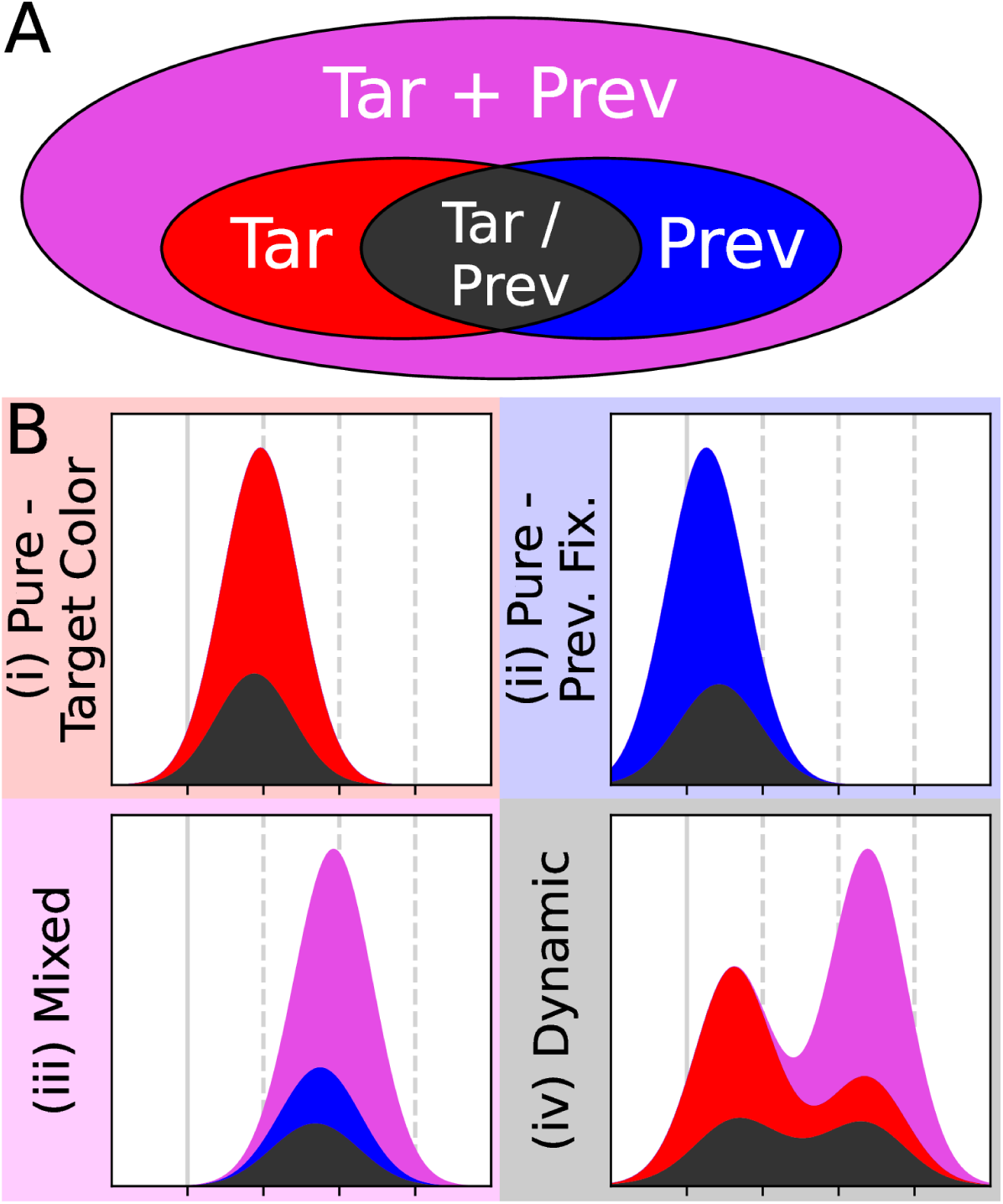
By creating distinct information signatures for each form of selectivity, PID can determine which one(s) are used by SC neurons to integrate target color and previous fixation signals in a visual search task. (A) The terms produced by a 2-variable PID analysis of the information encoded by a neuron’s firing rate: **redundant** information that can be predicted given access to either target similarity or previous fixation information (dark gray), **unique** information that can be predicted when and only when a model has access to target similarity information (red), **unique** information about previous fixation (blue), and **synergistic** information that can be predicted only given access to both the target similarity and previous fixation variables (magenta). This color scheme will be used throughout the paper. (B) Visualized hypotheses of different patterns of selectivity that neurons might exhibit. (i) Pure selectivity to target color would manifest in unique information about target color and possibly information redundant to target color and previous fixation. (ii) Pure selectivity to previous fixation would manifest in a corresponding pattern with unique information about previous fixation. (iii) Mixed selectivity would manifest primarily in synergistic information about both factors, perhaps with lesser amounts of unique information about one or both, and some redundant information. (iv) Dynamic selectivity might manifest as unique information about one term early in fixation, followed by synergistic information (relevant to stimulus value) later in target selection.

A strength of PID is that it enables each of these forms of information to be measured over time, and this is important because our research question requires us to consider the possibility that the information encoded by a neuron might change over time (i.e., dynamic selectivity). We therefore conducted a *time-series PID* analysis, which is a separate PID analysis performed at every millisecond. This allowed us to quantify how a neuron’s information selectivity changes over the course of saccade target selection and to determine what information a neuron is encoding at particular instants (e.g., fixation onset). We also introduce a novel filtering approach, discussed in Section 4.4, to limit the effects we report to information whose encoding is statistically significant in its temporal context.

PID has been successfully employed *in silico*, *in vitro*, and *in vivo*. In simulation, it has been used to understand how information flows through neural circuits performing minimal cognitive behaviors (Beer and Williams, 2015), including a short-term memory task (Candadai and Izquierdo, 2020). PID has also been used *in vitro* to quantify the effects of recurrence on synergistic neural integration in cortical cell cultures (Sherrill, Timme, Beggs, and Newman, 2021). In non-human animal experiments, PID has been used to quantify the establishment of new functional pathways during fear conditioning (Frontera et al., 2020), and a theoretically similar technique, Granger causality, has been used to quantify the information exchanged between two bats over the course of a vocalization (Rose, Styr, Schmid, Elie, and Yartsev, 2021). Overall, PID is a generally accepted method in the neuroscience literature (Timme and Lapish, 2018), if not yet one that is broadly used.

### 1.4 Aims and Hypotheses

To our knowledge, our study is the first to leverage the powerful PID method to investigate neural selectivity during attention control. In applying time-series PID, our aim is to determine which theory of neural selectivity best explains SC neurons’ encoding of the information necessary to control search behavior. In addition to this contribution to the theoretical neuroscience literature, another aim of our work is to further introduce the PID method to neuroscientists and to demonstrate its value in precisely indicating what information a neuron is encoding and when. Our hope is that our work will help neuroscientists better appreciate how PID might usefully complement other methods of quantifying neural selectivity.

To realize these aims, we formulate four specific hypotheses for how an SC neuron might encode the variables of target color and previous fixation to actively control eye movements in our sensorimotor task. These hypothesized patterns of neural selectivity are illustrated in Figure 2B and can be summarized as:

- Neurons found to encode unique information about target color will be interpreted as evidence supporting the purely selective encoding of the target-color factor.
- Neurons found to encode unique information about previous fixation will be interpreted as evidence supporting the purely selective encoding of the previous-fixation factor.
- Neurons found to encode unique information about both target color and previous fixation, or synergistic information about their interaction, will be interpreted as evidence broadly supporting mixed-selectivity theories in the context of SC.
- Neurons found to encode information about different factors at different times during the course of a fixation will be interpreted as evidence supporting dynamic-selectivity theories.

The SC is known to include multiple neuron types showing a diversity of responses (Sparks, 1978; Mays and Sparks, 1980; Gale and Murphy, 2014; Liu et al., 2022). PID has the potential to analyze these diverse responses to determine which type(s) of neural selectivity they represent. Motivated by work modeling SC’s population response as a priority map supporting target selection (Facteau and Munoz, 2006; Adeli, Vitu, and Zelinsky, 2017; Bisley and Mirpour, 2019), one potential finding is that SC neurons will predominantly focus their representational power on efficiently guiding saccades to unfixated potential targets based on expected reward – a variable which depends on both of our factors and so would be measured as mixed selectivity in our analysis. Alternatively, we may also find purely-selective neurons dedicated to inhibitory tagging, a computation which is fundamental to search and many other visual tasks. PID’s capacity to discriminate between these classes of neuron response highlights its value as a tool for studying neural selectivity in the primate SC.

## 2. Results

### 2.1 Diverse neural selectivity in SC

Figure 3 shows a striking diversity of neural selectivity patterns. Among the 113 neurons recorded by Conroy et al. (2023), we excluded 44 neurons that did not significantly encode either of the task variables. Of the remaining neurons engaged in the control of the grid-search task, we used a simple algorithm to assign PID signatures to selectivity theories (see Section 4.5) and confirmed this assignment by visual inspection. We found that 15 neurons showed pure selectivity to target color (Fig. 3A), evidenced by their predominantly red PID signatures, and another 17 neurons showed pure selectivity to previous fixation (Fig. 3B), evidenced in blue. We also found clear evidence for mixed and dynamic forms of neural selectivity. Mixed selectivity is visually classified as a representation of synergistic information (shown in magenta) or else two or more sources of information at the same time (temporally overlapping). In contrast, dynamic selectivity is visualized as the representation of different sources of information at different times (temporally distinct). We found that 15 neurons evidenced mixed selectivity (Fig. 3C), most commonly encoding information about previous fixation at the same time as synergistic information about both factors, and that another 22 neurons showed various forms of dynamic selectivity (Fig. 3D). Finding that the three forms of neural selectivity were comparably represented in this sample suggests that the attention control exerted by the SC broadly leverages multiple forms of neural selectivity.

**Figure 3.**
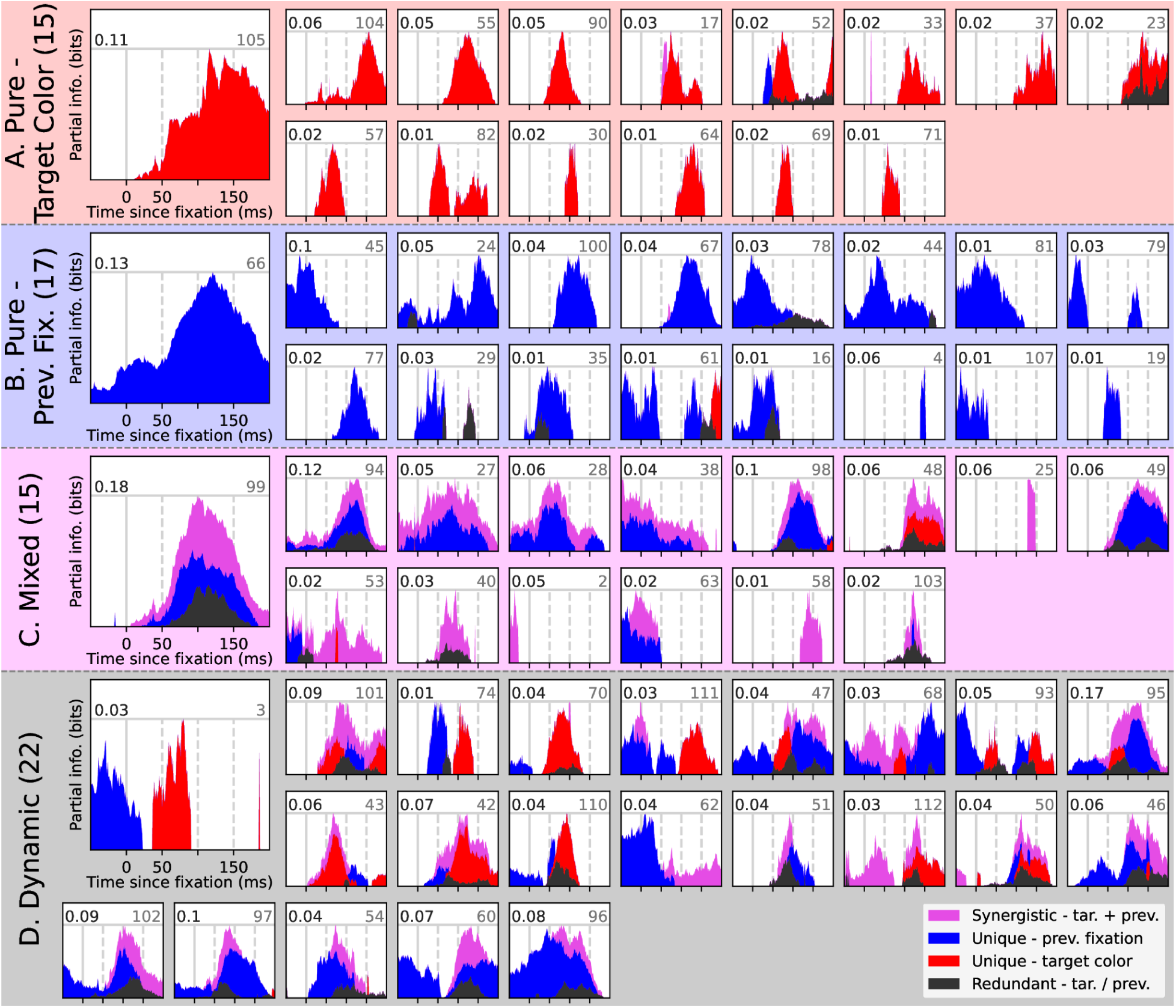
There is substantial variability in how SC neurons allocate their representational capacity over the course of saccade target selection. This figure shows time-series PID traces for 69 neurons, grouped into pure, mixed, and dynamic classes of neural selectivity (44 excluded). The information explained by each PID term is shown (stacked) on the y-axis. The maximum information explained by all terms together is shown with a solid gray horizontal line and written (in bits) at the top left of each panel. Each neuron’s index is written at the top right. Panels A-C are sorted by the quantity of information encoded about the relevant terms; panel D is sorted by the temporal separation of the relevant terms.

### 2.2 Previous fixation encoded first

In addition to PID’s demonstrated ability to categorize the use of different forms of neural selectivity by different neurons, we also exploited time-series PID’s temporal precision to ask when different types of information are encoded by neurons.

As can be seen by comparing Figures 4A and 4B, previous fixation was encoded substantially earlier than target color among purely selective neurons, with 12 of 17 previous fixation neurons encoding information about whether a disk was previously fixated while the saccade bringing this disk into the neuron’s RF was still in progress (<0ms before fixation onset). Only a single target color neuron (neuron 104) encoded any information this early, and the quantity of information was extremely small (refer to Figure 3A). In fact, only 5 of 15 encoded any significant target color information earlier than +50ms following fixation onset.

**Figure 4.**
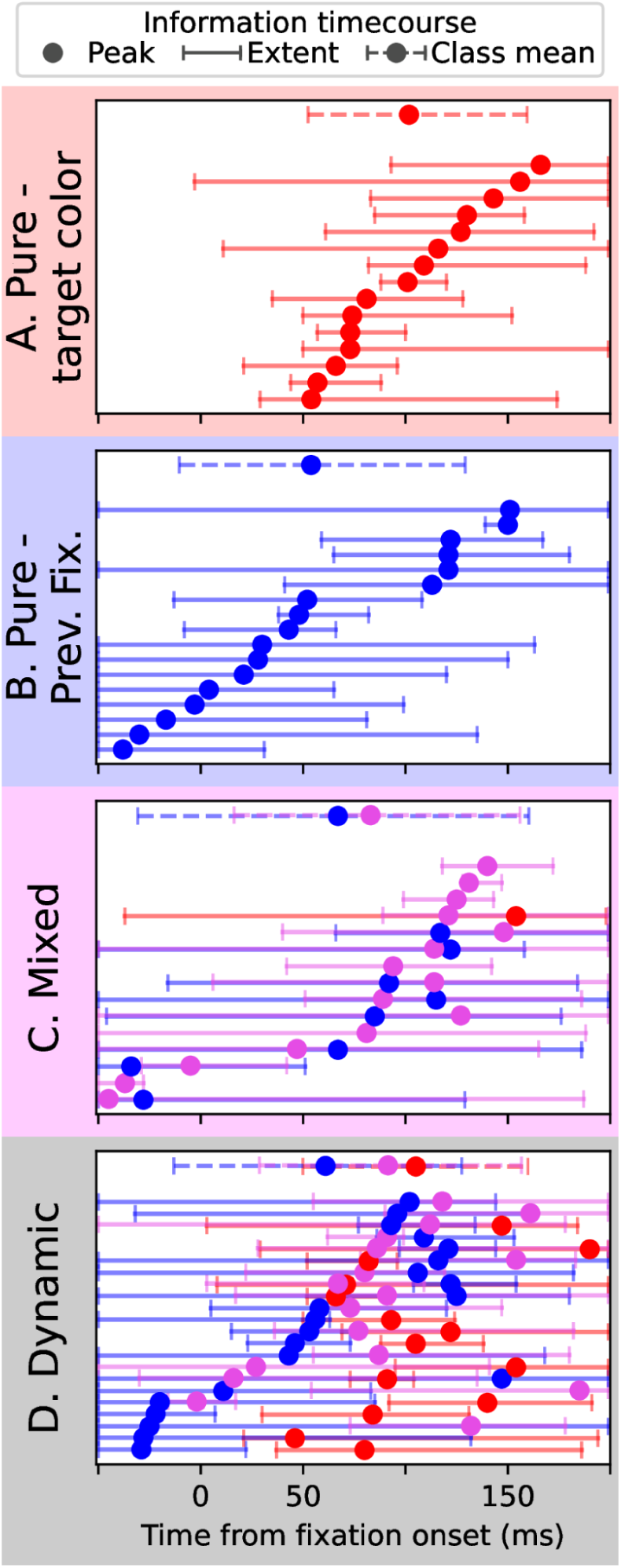
The information encoded by neurons of each selectivity class (A-D) changes over time. The y-axis of each panel indexes a count of neurons (shown individually in Figure 3), with each row corresponding to a different neuron. The top row in each panel (dashed line) is an average over all neurons in the class. Colors denote information terms, with unique information about target color match shown in red and about previous fixation shown in blue, and synergistic information shown in magenta. Circles code the moment that the neuron encodes its maximum information about a term; neurons are ordered based on this moment. Variability spans indicate the earliest and latest moments that the neuron encoded significant information about each term. See Section 4.4 for methods.

While there was some diversity among mixed-selective neurons (Figure 4C), their overall tendency can be approximated from the class average (dashed line). In general, this class of neurons encoded information about both previous fixation and its synergy with target color match over a prolonged period, with the synergistic information detected from +16ms to +156ms from fixation onset, on average.

Dynamically-selective neurons were even more diverse (Figure 4D), but they tended to encode unique information at times similar to the purely selective neurons, and synergistic information at times similar to the mixed selective neurons (compare average timecourses between the four panels). A typical response was to first encode previous fixation at fixation onset but then switch to encoding target color information after it becomes available from higher-level visual areas at +50ms following fixation onset (White and Munoz, 2011; Zhou and Desimone, 2011; Hafed et al., 2023). The 17 of 22 dynamic neurons that encoded synergistic information did so over a similar timeframe as the mixed neurons, with an onset and peak preceding those of target color. Time-series PID therefore reveals a diversity in both neural selectivity classifications and in the timecourses of information encoded by SC neurons.

### 2.3 Distributed over the visuomotor spectrum

The responses of SC neurons depend on their laminar depth: superficial-layer neurons are visually responsive, but lack saccade-related responses, while the responses of intermediate and deeper-layer neurons tend to become progressively more motor with increasing depth (Massot et al. 2019). To determine whether neural selectivity depends on this vision-movement dimension, we adopted the visuomotor index measure (VMI; Sparks, 1978), which is widely used in the contemporary study of SC (e.g., Shen and Paré, 2007; Massot et al., 2019; Caziot et al., 2025). This measure was available in our dataset; prior to the search task, Conroy et al. (2023) had computed each neuron’s VMI using a delayed-saccade task (see Section 4.1). We evaluated whether the neurons making up each neural selectivity class had a different distribution of VMIs compared to the distribution of the population overall. Figure 5 shows the results of this analysis. Using Mann-Whitney U tests, we did not find any significant difference in VMI for mixed-selective neurons (f = 0.46, p = 0.66), purely-selective target-color neurons (f = 0.53, p = 0.73), purely-selective previous-fixation neurons (f = 0.48, p = 0.76), or dynamically-selective neurons (f = 0.39, p = 0.12). Note that Conroy et al. (2023) recorded primarily from the superficial and intermediate layers of SC, and their dataset did not include any strongly motor neurons. It is therefore possible that such neurons would fall disproportionately in one of our neural selectivity categories, or be disproportionately excluded. However, among the predominantly visually-responsive neurons in this dataset, there is no clear or strong relationship between selectivity and VMI.

**Figure 5.**
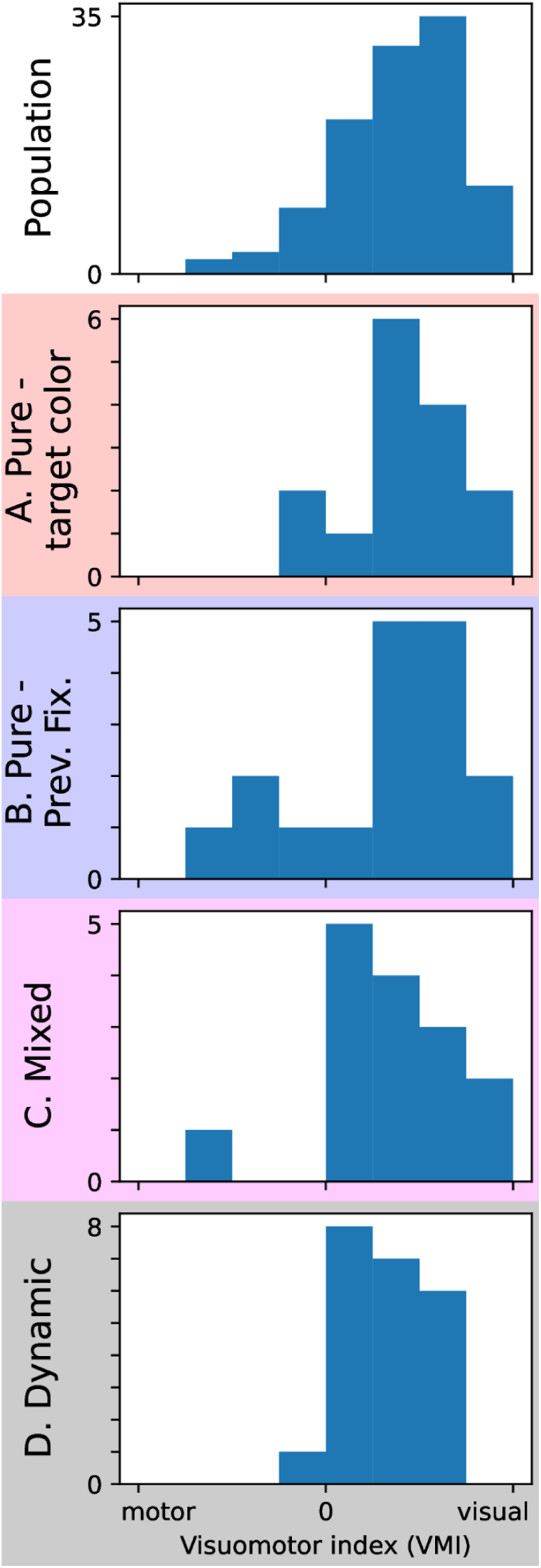
The distribution of visuomotor indices (VMI) within each neural-selectivity class (A-D) does not substantially differ from that of the whole population (top). Within each histogram, the y-axis indicates neuron count per VMI bin. VMI (shown on the x-axis) ranges from −1 (motor) to 1 (visual), with −1 indicating a neuron that responds exclusively to motor saccade planning and 1 indicating a neuron that responds exclusively to visual stimulation.

### 2.4 Validation with GLM

While PID does not rely on strong distributional assumptions and has clear advantages for interpretability, other techniques, in particular generalized linear models (GLMs), have been previously applied to questions of mixed selectivity. To ensure that the rich and interpretable findings from our PID analysis are consistent with the prevailing use of GLMs to assess neural selectivity patterns, we performed a parallel GLM analysis on our neural data (summarized in Figure 6) and compared its results with PID’s (Figure 3). Overall, we find largely similar patterns of results, with most differences stemming from standard GLM’s lower temporal precision and its normality assumption.

**Figure 6:**
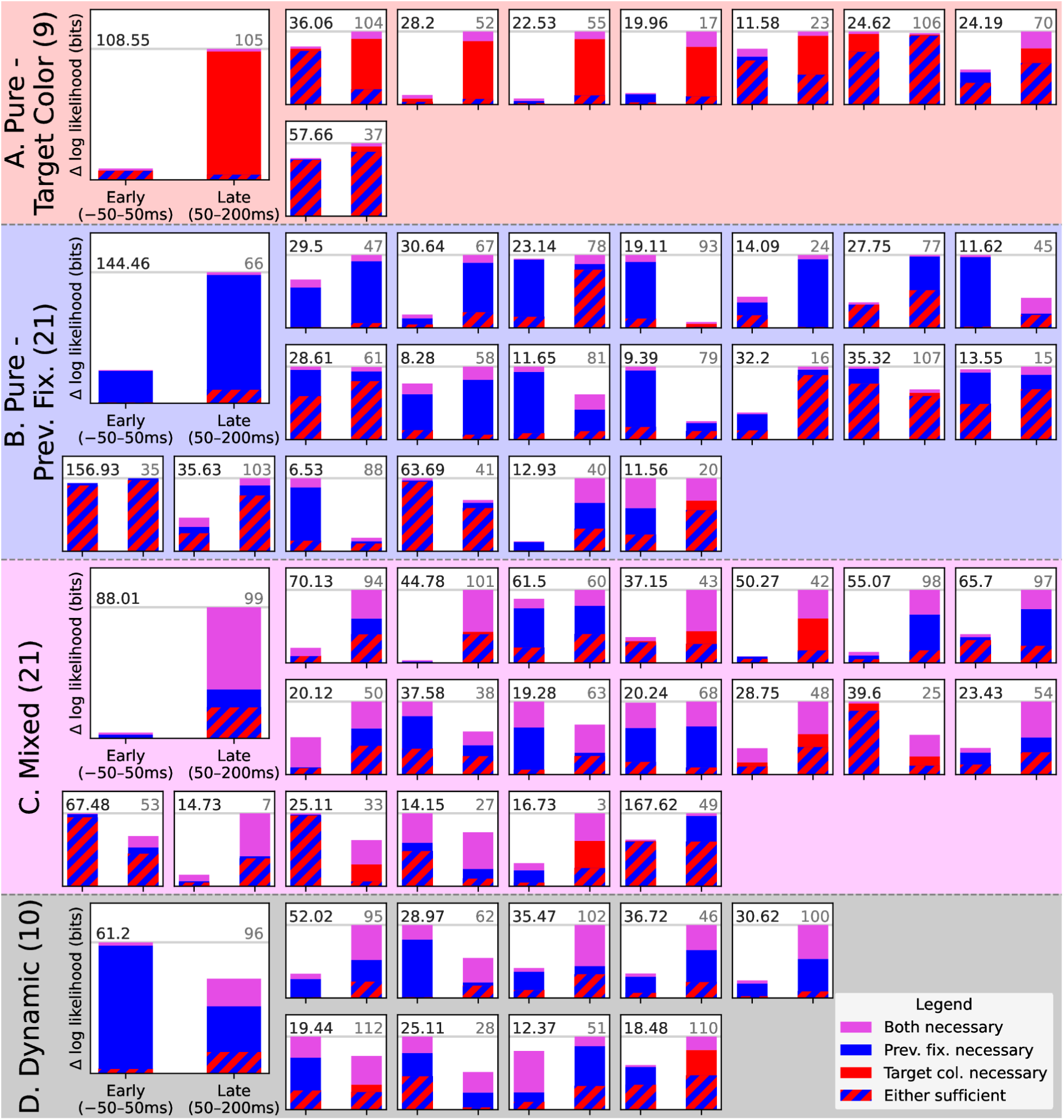
A complementary GLM analysis, computing explanatory power of linear models of pure selectivity incorporating single variables (red/blue), as well as a model of mixed selectivity incorporating their interaction (magenta). The stronger single-factor linear model is shown in a solid color; either factor is sufficient to explain the amount of information shown in a striped pattern. The y-axis measures the increase in log-likelihood vs a null model, with the greatest increase shown (in bits) at the top left of each panel. Neuron indexes are shown at the top right. Classifications of neural selectivity from this GLM analysis are computed separately from the PID classifications in Figure 3, but they largely agree (see Figure 7).

Our GLM analysis is identical to the single-factor GLM applied by Rigotti et al. (2013), and equivalent to the nested GLM approach employed by Diomedi et al. (2020), up to a difference in link function (see Section 4.6). Like those authors, we define mixing as the case where two variables are necessary to explain a neuron’s firing rate: this manifests in the model incorporating both factors having substantially higher likelihood than either of the single-factor models. We similarly define pure selectivity as the case where exactly one variable is necessary to explain the neuron’s firing rate, above and beyond the amount of (likely redundant) information that either variable is sufficient to explain. We label a neuron as exhibiting dynamic selectivity when its detected selectivity class differs during the early (−50ms to +50ms) and late (+50ms to +200ms) fixation periods.

The broad conclusions of this analysis are entirely consistent with our primary time-series PID analysis. Both analyses uncover substantial diversity in the selectivity patterns of SC neurons during search behavior, with distinct sets of neurons exhibiting pure selectivity to each task factor, mixed selectivity to them both, and dynamic selectivity that differs during early and late target selection.

Despite the broad agreement between the two methods, on a per-neuron level the GLM analysis does not always agree with time-series PID. Figure 7 presents a confusion matrix which shows that GLM and PID generally classify neurons the same way (66% of the time, with 5 classes), but which also highlights three common discrepancies between the analyses.

**Figure 7:**
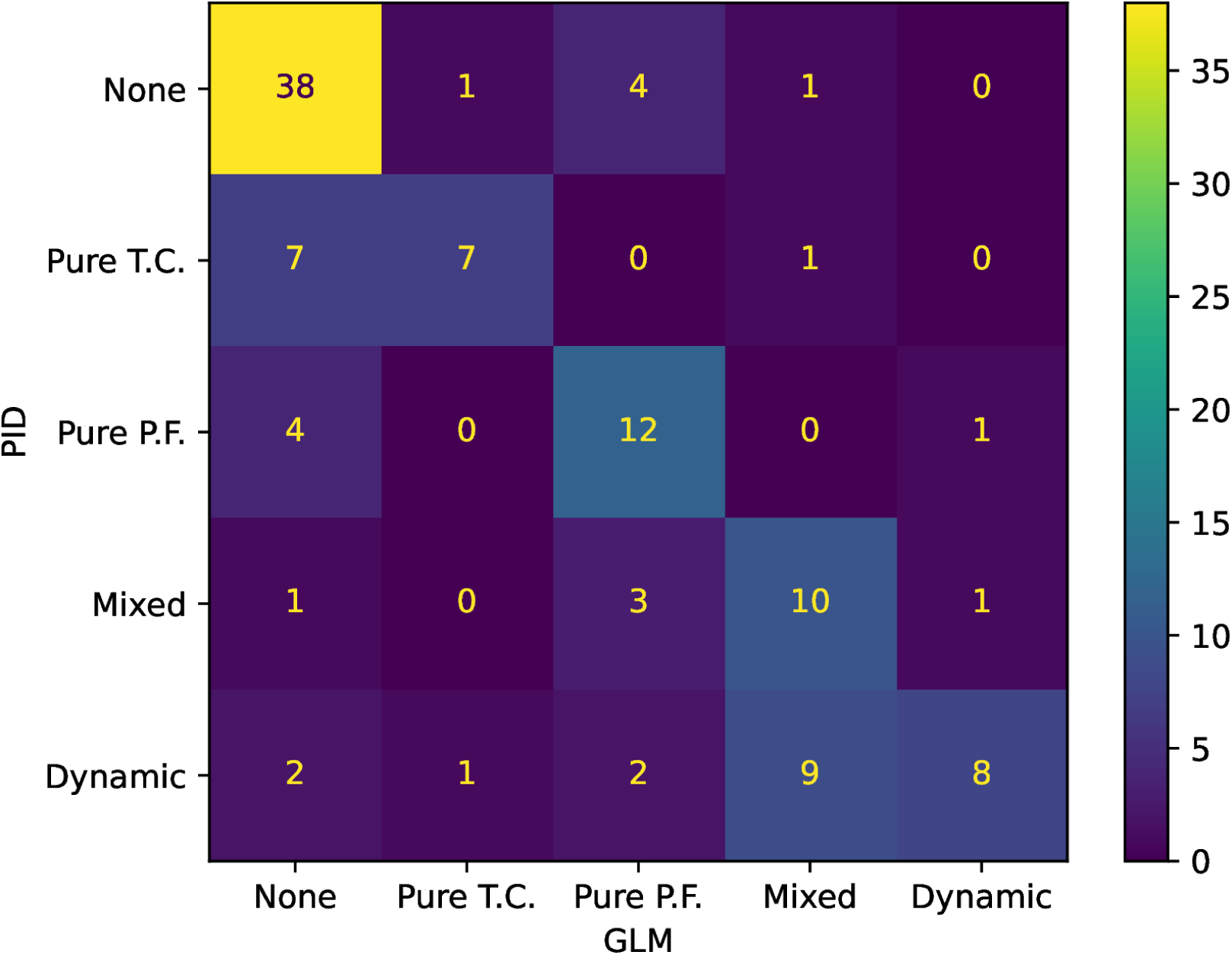
Although they generally agree, this confusion matrix enumerates cases where PID (y-axis, see Figure 3) and GLM (x-axis, see Figure 6) reached different conclusions regarding a neuron’s selectivity.

First, GLM reports that several neurons left unclassified by time-series PID (Figure 7, top row) exhibit pure selectivity to previous fixation (4 of 44). Of these neurons, GLM detected the strongest effect in neuron 15. As shown in Figure 8A, this neuron fires more for novel stimuli during the pre-fixation phase, though this effect is only marginally significant according to the parametric mean firing rate analysis shown in the left column. Due to our nonparametric Monte-Carlo significance filtering criterion, which does not assume that neural firing rates are normally distributed, the more conservative PID analysis reports that this effect is not significant. The more conservative analysis is appropriate here given that this neuron’s firing rates were not normally distributed (skew-kurtosis test, p<1e-5).

**Figure 8:**
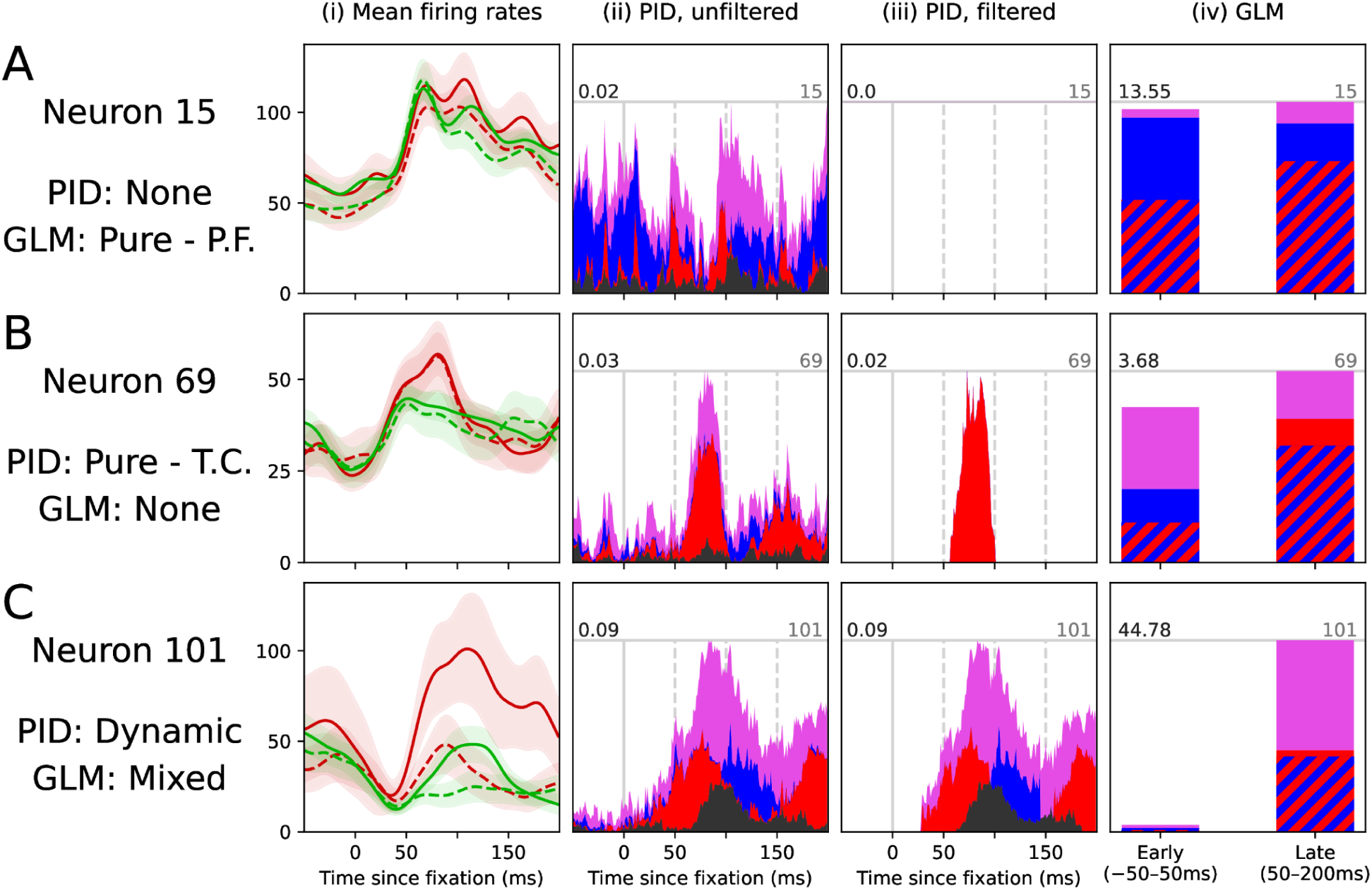
When PID and GLM disagree, we argue that PID provides the more accurate picture of a neuron’s role. This figure shows mean firing rates by condition (column i), unfiltered and filtered PID signatures (columns ii and iii), and GLM signatures (column iv) for 3 neurons where PID and GLM reported different classes. (A) Neuron 15 was not classified by PID, but GLM classified it as purely selective to previous fixation. (B) Neuron 69 was classified as purely selective to target guidance by PID and not classified by GLM. (C) Neuron 101 was classified as exhibiting dynamic selectivity by PID, and GLM classified it as exhibiting mixed selectivity. Color schemes follow the preceding figures.

Second, GLM fails to detect a sizable portion (7 of 15) of neurons where PID reports pure selectivity to target color. Neuron 69 (Fig. 8B) provides a striking example. GLM detects very little information about target color beyond that which is encoded about previous fixation (0.52 bits log-likelihood delta, Fig 8.B.iv), yet PID detects a clear and significant period during which it is encoded, +50-100ms after fixation onset (Fig 8.B.iii). Why the discrepancy? As it turns out, while this neuron used high firing rates during the +50-100ms window to indicate targets, it also had marginally *lower* firing rates for targets during the +150-200ms timeframe (Fig 8.B.i). Because standard GLM relies on averaging over time windows, it detects only a small difference in means between targets and distractors over the full +50-200ms window, and thus does not recognize this neuron’s brief but significant selectivity to targets.

Third, GLM assigns mixed selectivity as a label to a plurality (9 of 22) of the neurons where time-series PID detects dynamic selectivity. Focusing on one representative case, neuron 101 (Fig. 8c) exhibits distinct periods of encoding unique target color information, unique previous fixation information, and again target color information, with synergistic information being encoded over this full window (Fig. 8.B.iii). Because this neuron’s selectivity does not differ across the +50ms early-late divide, it is missed by the standard GLM analysis (Fig. 8.B.iv). Thus, if GLM had been the only analysis conducted, we would have understated the diversity of temporal selectivity patterns in the SC.

## 3. Discussion

### 3.1 PID can distinguish between patterns of neural selectivity

Ours is the first application of PID to the question of how neurons selectively encode the variables of an attention-guided task, and as such it contributes to the understanding of how attention control is implemented in the brain. We not only demonstrated that our novel time-series PID analysis can distinguish between theoretically-predicted patterns of neural selectivity, but in rigorous comparison we also showed that it can do so better than traditional generalized linear modeling approaches. Time-series PID also produces intuitive visualizations of information signatures consistent with each neural selectivity theory, which we hope will generate debate and theory refinement. We also hope that neuroscientists will find PID to be a useful supplement to their existing methods for interpreting neural data, allowing them to compute precisely what information a neuron encodes based on its firing rate distribution. To realize this hope our future goal is to integrate our time-series PID into an easily used toolbox.

#### 3.1.1 It does so better than GLM

In this work we introduce a new tool – time-series PID – to the study of mixed selectivity, which has typically been approached using generalized linear models (GLM). We found three specific patterns of discrepancies between GLM and time-series PID that have important implications for theories of neural selectivity. In each case, we argue that PID is the more complete and reliable analysis. These three patterns of discrepancies reveal the advantages of both our novel, statistically conservative significance filtering approach and time-series PID’s exquisite temporal and informational precision.

The first pattern, GLM’s false positives, stems from the fact that GLM and related models rely on strong distributional assumptions which generally do not hold over an entire neural population. A standard GLM employing a linear link function and Gaussian kernel assumes that its data are normally distributed, but this assumption was violated by every neuron in our dataset (skew-kurtosis tests). For neural populations as diverse as SC, nonparametric analyses like our Monte-Carlo significance filtering strategy are the most appropriate.

The next two discrepant patterns, PID’s stronger detection of pure and dynamic selectivity, emerge because our conservative statistical tests permit us to confidently detect and report effects with extremely fine temporal precision, as narrow as 1ms, while controlling familywise error. This precision is necessary to fully characterize the temporal selectivity of a neuron. The differential responses of neurons do not generally extend over the entire time windows used by standard GLM analyses, and sometimes effects are not even encoded in the same direction over this window. Because PID is computed at a fine temporal resolution, it is robust to these issues.

Beyond these empirical considerations, we also note that quantifying mixed selectivity using nested GLM (as in Diomedi et al, 2020) rests on an assumption of independence. Nested GLM reports mixing when there are multiple variables that are necessary to explain a neuron’s firing rate, and it reports the lack of mixing when there is only a single necessary variable. While this is intuitive, it fails in the face of correlated factors. Consider a firing rate well-explained by a linear combination of three factors, two of which are highly correlated. In this situation, omitting either of the correlated variables will hardly decrease the GLM likelihood at all – they are redundant with each other – but the uncorrelated variable is necessary. This neuron will then appear not to exhibit mixing under nested GLM. (Single-factor GLM would report mixed selectivity in this case because no single factor is sufficient to explain all variability, but it cannot quantify which specific variables are encoded.) Because PID explicitly localizes redundant information to its constituent factors, it can distinguish the unique and redundant sources of information that explain this neuron, and thus accurately report that it exhibits mixed selectivity. In a supplementary 3-factor analysis decomposing target guidance into moderately correlated salience and color guidance factors, PID detects such redundant mixing in neuron 98 (see Fig. S1F).

In spite of these advantages, PID cannot fully replace linear modeling in the characterization of neural selectivity. Unlike some GLM approaches, PID cannot distinguish between linear and nonlinear mixed selectivity, limiting its ability to estimate the dimensionality of a neural population response. Additionally, a complete PID in more than three variables is extremely data-hungry. Nested GLM reports one term per variable, corresponding to the explanatory power lost when each variable is omitted from the model. In contrast, the number of terms in a complete PID follows the Dedekind series (Dedekind, 1897), which grows super-exponentially in the number of variables. Our 2-factor PID reports 4 terms, but a 3-factor PID reports 18 and a complete 4-factor PID reports 166. Larger complete PIDs are likely impossible, but even incomplete PID analyses in more than 4 variables would yield insight beyond that of nested GLM.

### 3.2 SC neurons show patterns consistent with multiple neural selectivity theories

We found groups of SC neurons adhering to patterns predicted by each of three neural selectivity theories. With respect to pure selectivity, we interpret the presence of neurons purely selective to target color and previous fixation as indicating that the SC contains or interfaces with distinct circuits responsible for computing these two task variables. Our findings of mixed and dynamic selectivity provide insight into how the SC helps integrate the pure signals relayed by these circuits.

SC neurons purely selective to target color are part of a circuit receiving attention-biased color information from elsewhere in the brain. This signal may originate in V4, which is known to exhibit color-based attentional enhancement (Motter, 1994) and to project to the SC (Gattass et al., 2014). Another possible source is the frontal eye fields (FEF), which exhibit feature-based firing rate enhancement during extended search at a latency consistent with our findings: +50ms from fixation onset (Zhou and Desimone, 2011).

On the other hand, neurons purely selective to previous fixation participate in a different circuit, which is responsible for computing a history of previously fixated locations. This circuit may also involve other visuomotor areas, such as FEF and the lateral intraparietal area (LIP), each of which has been shown to encode previous fixation information on a similar task (Mirpour, Arcizet, Ong, and Bisley, 2009; Mirpour, Bolandnazar, and Bisley, 2019). One clue to the dynamics of this circuit is the relative latency at which different brain areas encode previous fixation. We found that SC encodes previous fixation prior to fixation onset, as does FEF (Mirpour et al., 2019); LIP, however, does not. This suggests that the previous fixation signal may originate in either SC or FEF, or may be computed jointly by the two areas.

In the context of the search task, the subpopulation of SC neurons exhibiting mixed selectivity controls efficient search by guiding attention toward disks having the target color and away from previous fixated disks. This task-dependent integration of information enhances the particular combination of variables that is most likely to yield reward for this task. While it is plausible that this integration occurs locally in the SC (given our evidence that its constituent inputs are available there), it may also occur elsewhere (e.g., FEF), or the computation may be distributed across multiple areas.

We speculate that the subpopulation exhibiting dynamic selectivity acts as a more local circuit, directly accumulating evidence to support saccade target selection. It is plausible that dynamically-selective neurons are driven by their more specialized neighbors in SC, because the dynamic neurons encode target color and previous fixation at roughly the same times as the mixed or purely-selective neurons dedicated to these variables. This hypothesis is consistent with existing work demonstrating locally excitatory dynamics in the intermediate SC (McIlwain, 1982; Pettit, Helm, Lee, Augustine, and Hall 1999; Lee and Hall, 2006; Phongphanphanee, Marino, Kaneda, Yanagawa, Munoz, and Isa, 2014; reviewed in Veale, Hafed, and Yoshida, 2017). Of the four subpopulations we discovered, this is the most mechanistically consistent with the view of SC performing a winner-take-all target selection on a priority map.

### 3.3 The SC is more than just a priority map

The prevailing model of the SC’s role in visual cognition is that it comprises a *priority map* that combines bottom-up information (received either from cortex or superficial SC) and top-down attention biases (Fecteau and Munoz, 2006; Zelinsky and Bisley, 2015; Adeli, Vitu, and Zelinsky, 2017; White, Berg et al., 2017; White, Kan et al. 2017; Bisley and Mirpour, 2019). Under this model, the function of the intermediate and deep layers of the SC is to perform local winner-take-all (or probabilistic [Basso and Kim, 2010]) computations to select a single saccade target on this map (e.g., Trappenberg et al., 2001; McPeek and Keller, 2004; Carello and Krauzlis, 2004; Bisley and Mirpour, 2019). Even the study motivating the present work, Conroy et al. (2023), frames its findings through the lens of the SC encoding “a priority map used for saccade target selection” (p. 824).

Our findings clearly indicate that this “priority map” view of the SC is incomplete. At the very least, SC participates in specialized circuits that encode more specific information than what is required for target selection from a priority map. SC is generally understood to receive a top-down attentional bias from cortical areas in pre-integrated form. Under this view neurons purely selective to the factors contributing to this bias should not exist in SC, but we found them. What’s more, we found evidence for dynamically-selective neurons and identified a plausible mechanism by which they might combine the signals encoded by purely-selective neurons. Taken together, our findings offer compelling evidence that the SC is not exclusively dedicated to priority, and we argue by parsimony that it likely performs some task-relevant computations locally.

We are not alone in this conclusion. The SC has been shown to support object perception in acortical mice (K. H. Lee, Tran, Turan, and Meister, 2020), and it has recently been shown to enhance face stimuli with latencies as low as 40ms using signals routed through LGN (Yu, Katz, Quaia, Messinger, and Krauzlis, 2024). In a recent review, Cruz et al. (2023) argue that the SC performs computations locally, including the salience coding identified by White, Kan, et al. (2017), and then relays the results to cortical areas such as the thalamus (specifically the pulvinar and LGN) (Ahmadlou, Zweifel, and Heimel, 2018). Finally, primate SC was shown to support the recognition of learned novel categories in a task not requiring eye movements (Peysakhovich et al., 2024).

Our work joins this growing chorus of research in arguing that SC performs sophisticated, task-dependent computations to actively control visual search behavior. Using a novel information-theoretic analysis, we found evidence for pure, mixed, and dynamic selectivity in SC. We see this as convincing evidence that the SC is not only a structure for selecting a target from a priority map, but also the structure that creates this priority map by allocating the neural resources and local computations needed to integrate multiple sources of top-down task information.

### 3.4 Limitations and future directions

Our work provided compelling quantitative evidence for the existence of discrete neural circuits in the SC controlling efficient search behavior. However, our work leaves two theoretically important questions unanswered. First, it remains uncertain whether the purely-selective neurons we discovered are dedicated to their computations across all behavioral contexts, or whether they are task-dependent. Addressing this question would require data collected from the same neurons over multiple tasks, as Yang et al. (2019) did in simulation, and in future work we hope to take a first step toward answering this question by collecting grid-search data and guided-saccade data from the same neurons. Second, the nature of the mixed selectivity that we discovered in SC is unclear. It may be primarily task-variable selectivity encoding the expected reward associated with a saccade, as predicted by priority map theory, or it could be primarily category-free selectivity encoding arbitrary combinations of stimulus properties, including those irrelevant to a task. Given that our current experimental design included only two variables, both of which were task-relevant, our present analysis is incapable of distinguishing between these cases. In future work we hope to add a luminance-salience variable, and use a three-variable PID (akin to the one shown in Supplemental Figure S1) to answer this remaining theoretical question about neural selectivity in the SC.

## 4. Methods

### 4.1 Neural data

We provide a brief summary of the behavioral and neural data collection procedure from Conroy et al. (2023). Microelectrodes were used to record isolated single units from the superior colliculi of two male rhesus monkeys (*macaca mulatta*) as they performed the delayed saccade and search tasks described below. Eye movements were recorded using a video-based eye tracker (Eyelink 1000, SR Research). Voltages and concurrent eye-movement data (saccade and fixation onsets) were recorded synchronously during each trial.

Prior to data collection for the multi-target grid search task, Conroy et al (2023) recorded each neuron’s response during a standard delayed saccade task in order to localize its RF and to compute its visuomotor index (VMI). In this task, monkeys held central fixation on a disk stimulus for 450-650ms, after which an additional disk appeared in the periphery. Monkeys were required to maintain fixation on the central disk until it disappeared (following an additional 500-700ms), at which point an eye movement was made to the peripheral disk. The neuron’s RF was localized by manually varying the peripheral disk’s location to maximize visual and motor responses. Following localization, the neuron’s VMI was computed by averaging its response over at least 10 more trials. VMI followed the formula 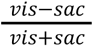, where *vis* is the neuron’s mean firing rate during the 50ms-100ms period following the onset of the peripheral disk and *sac* is the neuron’s mean firing rate during the −50ms-0ms period preceding the onset of the saccade.

Our analyses focused on the data recorded during the search task. At the beginning of each trial, the monkey was required to hold fixation on a central disk (not sharing the target color) for 500ms. Following this fixation, a grid search array comprising up to 24 additional disks (25 total) was shown. These disks differed only in color; one third of the additional disks shared the target color (either red or green, varying by trial block). This array was rotated and scaled such that the recorded neuron’s receptive field would fall on a disk near the currently-fixated disk; for highly eccentric RFs this required the array to include fewer than 24 disks. Only one of the disks was the true target, upon whose fixation the monkey received liquid reward and the trial ended. Trials would also end following a timeout of 8s. On roughly 7% of trials, every disk sharing the target color was a true target yielding reward upon its first fixation, and the trial terminated only once all targets had been fixated (or the trial timed out); Conroy et al. describe this condition as motivational.

The neural spike data were smoothed with a Gaussian kernel (σ = 10*ms*) to obtain smoothed firing rate estimates. Firing rate analyses targeted a −50ms to +200ms window around each fixation onset. The neural data were labeled for the properties of the visual stimulus in the recorded neuron’s RF during each fixation (target or non-target color), whether the stimulus had been fixated earlier in the search trial, and whether the saccades preceding and succeeding the fixation were into the neuron’s RF. To control for visual and motor factors confounding our results, our analyses excluded from this dataset cases in which fixations resulted in no stimulus falling in the recorded neuron’s RF and cases in which the prior or next saccade was into the RF of the neuron.

### 4.2 Partial information decomposition

We performed partial information decomposition using the standard methods introduced by Williams and Beer (2010). Partial information decomposition is an information theoretic analysis whose terms are constructed from quantities such as *entropy* (the amount of information needed to specify a variable) and *mutual information* (the amount that knowing one variable reduces the entropy of another – which happens to be symmetric). Concretely, PID partitions the mutual information between a target variable (such as neural firing rate) and the joint distribution of multiple source variables (such as the two factors in our experiment) into terms representing the redundant, unique, and synergistic combinations of these variables. These combinations of variables form a partially ordered set or lattice linked by containment: for instance, the term corresponding to the synergistic interaction of two variables contains each of the terms corresponding to a single variable’s unique information; those unique terms in turn contain the term corresponding to the two variables’ redundant interaction.

A PID lattice is computed from bottom (most redundant) to top (most synergistic). For each term in the lattice (representing a particular redundant or synergistic combination of source variables), one first directly computes the *total information* of the term – the extent to which the variables explain the target variable. From this, one can obtain the *partial information* associated with the term by subtracting all lower (contained) partial information terms in the PID lattice from the total information of the term in question. For the lowest term in the lattice (the redundant interaction of all source variables), there are no lower terms, so the total and partial information are the same.

The total information of unique and synergistic combinations of source variables is defined as the mutual information between the joint distribution of source variables and the target variable. Since mutual information is a well-known measure, the primary challenge becomes defining the total information that a redundant combination of source variables provides about the target variable.

Williams and Beer’s answer is *I_min_*, which reports the extent to which any given response provides information about multiple source variables at once. It is computed by taking the expected outcome of the following three-step procedure. First, sample a value *s* of the target variable *S* (neural firing rate). Second, compute how much knowing that *S = s* reduces the entropy of each source variable *A_i_* – in Williams and Beer’s notation, this is the specific information *I A_i_*; *S* = *s*. Finally, given a sampled *s* and the specific information that *S = s* gives us about each source variable *A_i_*, report the *lowest* specific information value across all source variables *A_i_*. This quantity represents the redundant re-use of a particular firing rate (eg, zero firing rate) to encode information about multiple factors. *I_min_* is defined as the expectation of this minimum over samples *s ∼ S*; that is, 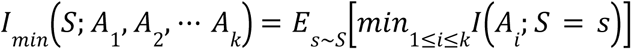.

### 4.3 Time-series PID

In order to apply PID to our time-series data, we performed separate PID analyses at each timestep. We applied PID to the smoothed firing rate data, which we found yielded more interpretable results than applying PID to spike counts (even over moderately large time bins of 10ms). This is theoretically justified given observed variability in response timing. The PID analyses at each timestep are correlated with those at neighboring timesteps due to both natural temporal continuity and smoothing. This made significance testing challenging, and we needed to develop new techniques, described in the following subsection, to perform it.

A second challenge in applying PID to neural firing rate data is to turn the continuous variable of neural firing rate into a discrete variable. We did so by binning firing rates into their instantaneous quartiles. That is, the bin boundaries were calculated separately for each neuron and each timestep. The choice to use equal-size quartile bins in every PID analysis was motivated by a desire to make all reported PID terms comparable, and it determined the neuron’s represented maximum information capacity, which was 2 bits. Note that when a neuron did not fire for more than 25% of fixations, the bottom quartile would include all such fixations, so that the neuron’s actual represented information capacity would be lower than 2 bits. While the decision to use equal-size bins was theoretically motivated, the decision to use quartiles in particular was driven by the observation that the neuron’s information explainable by the task factors plateaued at this point, but the information explainable by the null model (see below) continued to grow, indicating a noise ceiling.

### 4.4 Time-series significance filtering

Unlike the other analysis tools we used, our time-series significance filtering method is an entirely new contribution. We therefore describe it in detail.

Our significance filtering is based on Monte-Carlo and closed testing principles. We used a null model of each neuron’s data that preserved the neuron’s distribution of firing rates at each moment and also preserved the distribution of stimulus properties, but broke any relationship between the two. We did so by randomly shuffling the neuron’s firing rate over fixation indices, while keeping the original stimulus properties for each fixation. We then performed full PID analyses for each Monte Carlo run, yielding the same four terms reported in our primary analysis, to understand how much ‘information’ the neuron would have encoded about the stimulus properties in a distribution with no true mutual information between the two.

Given this null model, our time-series significance filtering can be conceptualized as occurring over two steps: restriction and expansion. The first step is to restrict reported results to cases where we are confident that a neuron encodes a particular term at a particular instant. To do this, we obtain a significance threshold for each neuron and term corresponding to a 95% confident judgment that a response of this magnitude would not have occurred *at any time* during a randomly sampled Monte Carlo run. Concretely, we compute the maximum over time of each PID term for each Monte Carlo run, and then take the 95th-percentile value over runs as our threshold. In the restriction step, we tentatively exclude all PID measures which do not surpass this threshold. This procedure is sound under closed testing principles and strikes a balance between a naïve 250-way Bonferroni multiple tests correction (which would be overly conservative given the time-correlated nature of our data) and an uncorrected analysis (which would be far too liberal and fail to prevent potential spurious results).

Having determined that a neuron encodes a particular term at a particular instant, in a second step we expand the duration of the reportable coding period. Expansion starts with the 1-timestep windows that restriction found to be significant, and then extends each window in each direction until neither the start nor the end of the window can be extended with 95% confidence. Interpreting an extended period of neural response as an accumulation of evidence that this response is genuine, we threshold the *n*th timestep of a response period, not on the 95th percentile of the maximum single timestep, but rather on the 95th percentile of the difference in accumulated information between the maximum response over a period of *n* − 1 timesteps and the maximum response over a period of *n* timesteps. For example, a significant period of coding from +30ms to +34ms may be extended to +35ms only if the information term at 35ms exceeds the 95th percentile (over Monte Carlo runs) of the ‘best 5th millisecond’ – the difference in summed information between the most informative 4ms window and the most informative 5ms window.

All primary analyses were based on 5000 Monte Carlo runs; our supplementary 3-variable analysis used 1000 Monte Carlo runs. These significance tests were by far the most computationally expensive part of this paper, and took roughly 60 minutes to complete on a 2021 MacBook Pro with an M1 Max CPU and 32 GB of RAM.

We encourage those interested in reproducing this significance filtering technique to refer to Supplemental Figure S2 for detailed visualizations of its intermediate steps on our dataset.

### 4.5 Neuron classification

We classified neurons to forms of selectivity using an algorithm that was developed based on visual inspection of the PID plots. This algorithm operates on each neuron by determining first which PID terms the neuron encodes substantial information about, and second whether these PID terms are encoded at substantially different times.

In determining the presence of substantial information, we aim to exclude cases where a neuron allocates a very small portion of its representational capacity to incidentally encode some extraneous variable. These cases are generally transient and are not visible when inspecting firing rate distributions; they were excluded using a threshold of 0. 25*bit* · *ms* on the area in the PID plot where PID detects significant information about the variable. This threshold corresponds to a neuron encoding 0.025 bits of information about the term (an eightieth of its represented information capacity) for a period of 10ms. Sub-threshold terms were encoded by neurons 17, 52, 53, 61, 67, and 98 and are visible as very small areas of color in Figure 3.

To determine whether a neuron encodes different terms at different times, we use the intersection-over-union metric on the timesteps when the neuron significantly encodes each term. To calculate this, we divide the number of milliseconds when the neuron encodes both terms (stacked in the PID plot) by the number of milliseconds when the neuron encodes either term. We consider encoding to occur at distinct times when the intersection-over-union of two terms is below 50%. As is visible in Figure 3, neuron 96’s intersection-over-union of unique information about previous fixation and synergistic information is just below 50%, whereas neuron 63’s intersection-over-union of these terms is above this threshold, leading to their different classifications.

Having this information about each neuron, we classified its neural selectivity using the following criteria:

(1) a neuron that encodes unique information only about previous fixation, and does not encode any synergistic information, was labeled as purely selective to previous fixation.
(2) a neuron that encodes unique information only about target color, and does not encode any synergistic information, was labeled as purely selective to target color.
(3) a neuron that encodes unique information about multiple factors, or else encodes synergistic information, but which does not encode different terms at different times, was labeled as exhibiting mixed selectivity.
(4) a neuron that encodes information about multiple terms at different times was labeled as exhibiting dynamic selectivity.
(5) a neuron that fits none of these selectivity patterns did not receive a classification and was excluded from our plots and downstream analyses.

### 4.6 GLM analysis

We performed a GLM analysis on our data primarily as a comparison to PID to help readers (who are likely to be more familiar with GLM) build an intuition about PID. We thus kept this analysis as simple as possible.

We used the Python statsmodels toolkit (Seabold and Perktold, 2010) to facilitate this analysis. Following general practice, we split time-series firing rate into an early (−50ms to +50ms) and a late (+50ms to +200ms) window relative to fixation onset. The window boundary was selected based on the general consensus that incoming color information is broadly available in SC at this time (e.g., White and Munoz, 2011; Zhou and Desimone, 2011), and the boundary is reasonable given our findings that neurons with dynamic selectivity tend to begin encoding target color match information at this time (see Section 2.2). Thus, the early window primarily captures internal processing, and the late window primarily captures stimulus-driven and external processing.

We fit four models for each neuron at each time window: (i) a mixed selectivity model including previous fixation, target color match, and their interaction, (ii) a pure model including only previous fixation, (iii) a pure model including only target color match, and (iv) a null model including no variables.

Following Rigotti et al. (2013), our GLMs used a linear link function and a Gaussian kernel, meaning that neural firing rate is assumed to vary linearly with neural stimulation (i.e., incoming firing rates), with a Gaussian (normal) error distribution. For these models, this is equivalent to describing a neuron’s firing rate by its mean and standard deviation under each combination of the variables included in the model. (Because there is an interaction term in our mixed model, it detects both linear and nonlinear mixed selectivity.) We used two-fold cross-validation. Our analyses yield four model likelihoods at each time window, where each likelihood refers to the probability of obtaining the observed data sample if the model were the true distribution. We take the log base 2 of each likelihood and subtract the log likelihood of the null model to obtain the information gain (in bits) of each model relative to the null model.

Following manual comparison of each neuron’s GLM likelihoods to its firing rate data, we pragmatically derived the following algorithm to classify each neuron’s selectivity at each time window: (a) if the log likelihood of the mixed model is more than 6 bits greater than the stronger pure model, label the neuron as mixed; (b) if the log likelihood of the pure previous fixation model is more than 4 bits greater than the pure target color guidance model, label the neuron as pure previous fixation; (c) if the log likelihood of the pure target color model is more than 4 bits greater than the pure previous fixation model, label the neuron as pure target color; (d) fail to label the neuron at this time window.

This algorithm results at two labels for each neuron, corresponding to its early and late selectivity. The labels at the two time windows are combined as such: (a) if the windows have different labels, label the neuron as dynamic; (b) if both windows have the same label, or only one window has a label, label the neuron with the single label; (c) fail to label the neuron.

## Acknowledgements

This paper is based upon work supported by the National Institutes of Health under Grant No. R01EY030669 and work supported by the National Science Foundation Graduate Research Fellowship under Grant No. 2234683.

---

**Supplemental Figure S1.**
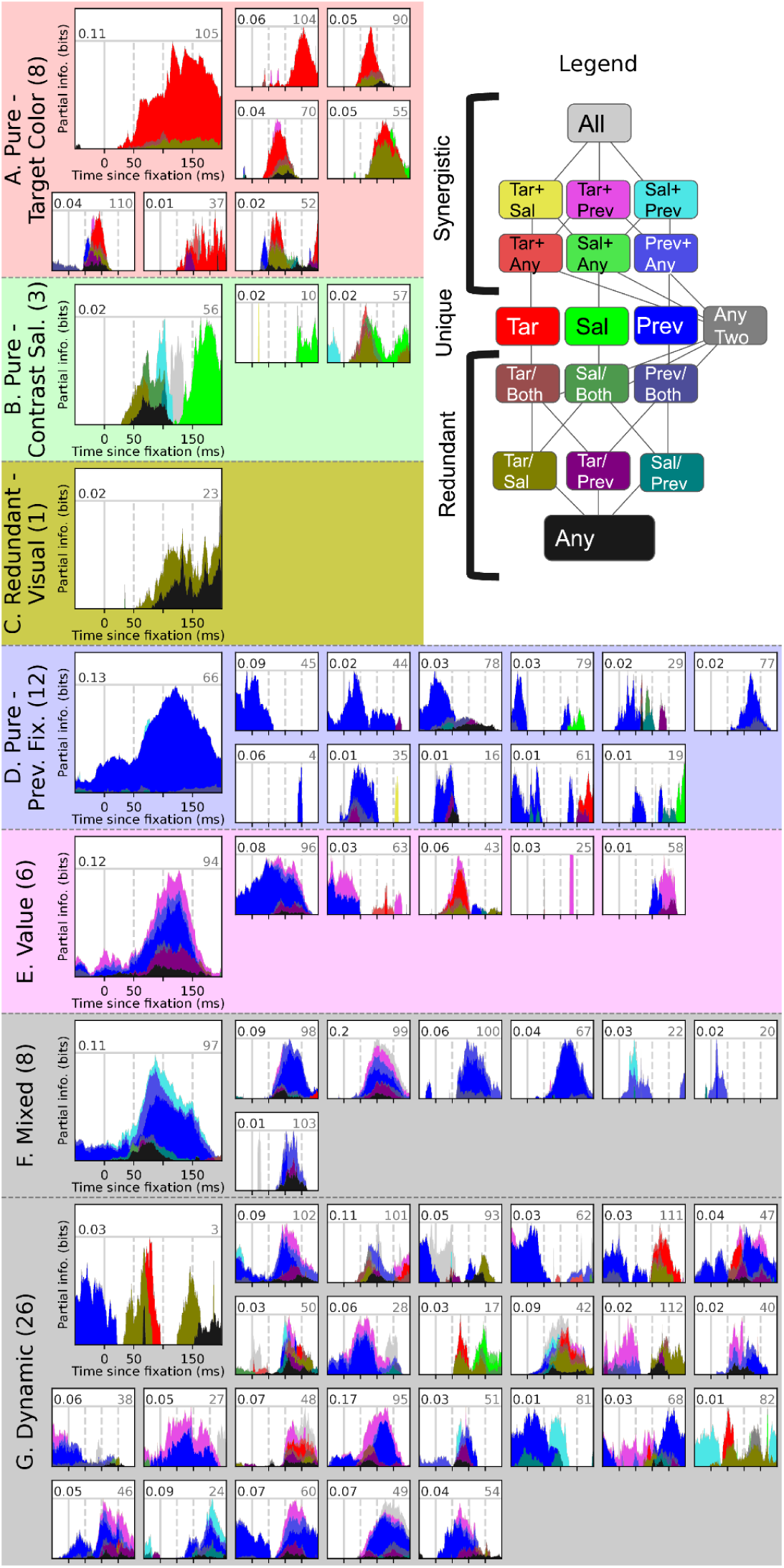
A 3-factor PID analysis on this neural dataset, partitioning visual information into color and contrast guidance signals and studying their relationship with previous fixation information. The legend is color-coded using RGB channel mixing with red denoting target color, green denoting contrast salience, and blue denoting previous fixation. Redundant interactions are denoted using ‘/’ and dimmed colors, and synergistic interactions are denoted using ‘+’ and bright colors. Figure design follows Fig 3.

## S1. 3-factor PID reveals stronger coding of target color than contrast salience

Because there were fewer potential targets (33%) than distractors (66%) in Conroy et al (2023)’s search arrays, we considered the possibility that target guidance resulted from an odd-one-out color contrast salience signal as well as the color-based attentional bias. That is, on a trial where potential targets were shown in red, red disks were more likely to be surrounded by green disks than vice versa. Thus, color-based guidance to the red targets was correlated with a potential odd-one-out salience signal. We performed an exploratory analysis to decompose target guidance into these target color and contrast salience signals.

A 3-factor PID finds that visual selectivity in the superior colliculus on this task is primarily dedicated to target color information, even when considering odd-one-out contrast salience. Overall, we found little evidence for the contrast salience signal we had hypothesized and thus confine our discussion of the signal to this supplementary analysis.

PID is robust to correlated predictors, and reports neurons that cannot disambiguate between predictors as encoding redundant information. 15 neurons supported target guidance in the primary 2-factor PID, and our supplementary 3-factor PID classified them as follows:

- 6 were purely selective to target color (Fig S1A, neurons 37, 52, 55, 90, 104, and 105).
- 1 redundantly encoded target color and contrast salience early during fixation and purely encoded contrast salience during late fixation (Fig S1B, neuron 57).
- 1 redundantly encoded visual information that could not be localized to either visual factor (Fig S1C, neuron 23).
- 2 dynamically encoded target color and a second later term involving contrast salience (Fig S1G, neurons 17 and 82).
- 5 were no longer found to significantly encode target color due to the higher statistical power required by 3-factor PID (not shown in Fig S1).

The 3-factor PID uncovered 3 neurons in total that primarily encoded contrast salience (Fig S1C), and 8 neurons (Fig S1F) that encoded synergistic information but could not be established as specifically encoding the synergy between target color and previous fixation (shown in magenta) that represents value on this task. In most cases, this was because those neurons encoded information that could be predicted by the synergy between previous fixation and (redundantly) either target color or contrast salience (‘Prev+Any’, shown in light blue). One neuron (Fig S1F, neuron 99) significantly encoded a synergy between all factors (shown in light gray). We describe those 8 neurons as exhibiting an unspecified version of mixed selectivity and contrast them with the 6 neurons (Fig S1E) that specifically encode expected value on this task (refer to Hirokawa et al, 2019, for a possible interpretation of this contrast). In future work, we plan to more thoroughly compare value selectivity and mixed selectivity in SC using a planned luminance salience manipulation, rather than post-hoc contrast salience.

While we found little evidence for a contrast salience signal, we draw three indirect conclusions from this supplementary analysis: (1) SC robustly encodes target color information when controlling for odd-one-out contrast salience; (2) a 3-factor time-series PID, yielding 18 terms, is feasible and yields insight even given the limited number of trials afforded by single-unit neurophysiology; but (3) the 3-factor PID will exhibit somewhat more type II errors (false negatives) given its higher power requirements.

### S1.1 Methods

Contrast salience was operationalized by labeling a disk as salient when its color differed from more than half of its neighbors’ colors (out of up to 4 immediate neighbors and up to 4 diagonal neighbors, weighted by 50%).

Neurons were classified using an algorithm identical to our primary analysis, with two changes: (i) neurons encoding unique information about one visual factor as well as redundant information about both visual factors were labeled as purely selective to the visual factor they encoded unique information about; (ii) neurons encoding synergistic information about target color and previous fixation, and no other non-redundant 2- or 3-way synergies, were labeled as exhibiting selectivity to expected value.

**Supplemental Figure S2.**
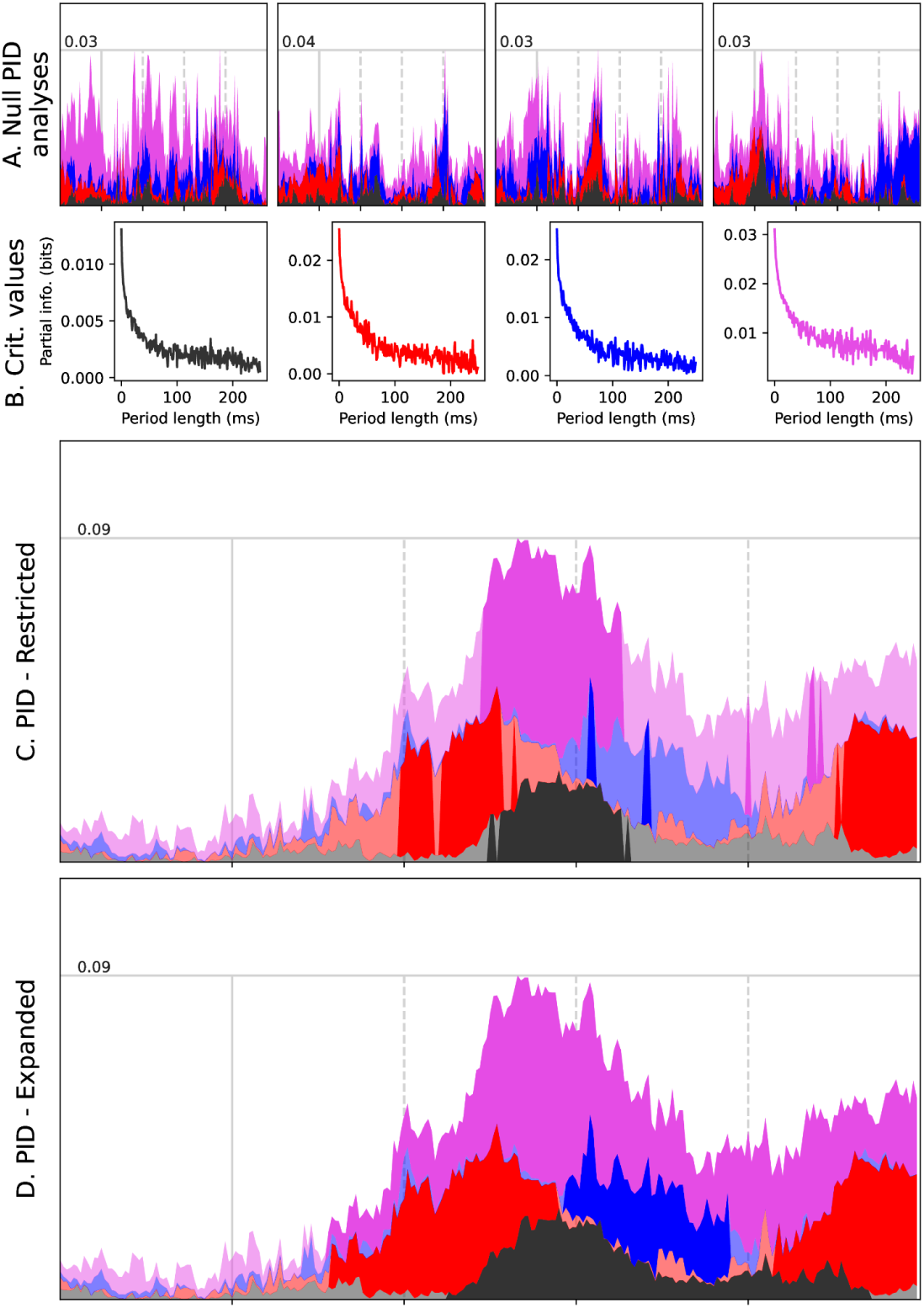
The intermediate steps of our novel time-series significance filtering method (described at length in Section 4.4), as applied to neuron 101. (A) Four of the 5000 Monte Carlo runs. In each run, stimulus properties and neural responses were dissociated and shuffled to act as a null model, and then PID was performed on the resulting shuffled data. The resulting signatures are spiky but include some clustering, resulting from the temporally-correlated nature of the input data. (B) Critical value lookup curves for each PID term, derived from 95th-percentile values over the set of Monte Carlo runs. These are used for both restriction and expansion. The x-axis shows the length of the window considered for expansion. The y-axis shows the quantity of information that must be exceeded at the edge of the window to allow expansion. (C) For restriction, the leftmost critical value, corresponding to the greatest amount of information encoded at any millisecond, is used as a threshold over the unfiltered PID time series. Information terms that did not exceed this threshold are faded out on the PID plot shown. (D) For expansion, each window that passed the restriction stage is expanded until the marginal information added by expanding the window another millisecond is less than the critical value at the window’s current length. Information terms that did not pass expansion are faded out on the PID plot shown here, and they are omitted from the filtered PID plots presented in the main text.

## Footnotes

1 Or, in nested modeling approaches to mixed selectivity (Diomedi, Vaccari, Filipini, Fattori, & Galetti, 2020), one tests whether multiple variables are *necessary*, meaning that a model omitting either of them is worse than a model incorporating all variables. These two approaches are equivalent in a two-variable analysis.

2 While unique and synergistic information are quite intuitive, redundant information is a bit more subtle. Properly, under the standard articulation of PID, redundant information refers to cases where a neuron uses a single response to communicate information about multiple source variables. See Section 4.2 for details.

## Notes

### Competing Interest Statement

The authors have declared no competing interest.

